# Mantis shrimp rank the shape of an object over its color during recognition

**DOI:** 10.1101/2020.08.20.259903

**Authors:** Rickesh N. Patel, Veniamin Khil, Laylo Abdurahmonova, Holland Driscoll, Sarina Patel, Olivia Pettyjohn-Robin, Ahmad Shah, Tamar Goldwasser, Benjamin Sparklin, Thomas W. Cronin

**Affiliations:** University of Maryland, Baltimore County. UMBC Department of Biological Sciences, 1000 Hilltop Circle, Baltimore, Maryland 21250

**Keywords:** object recognition, learning, memory, ethology, Pavlovian conditioning, animal behavior, mantis shrimp, stomatopod, marine biology, visual guidance

## Abstract

Mantis shrimp are predatory crustaceans that commonly occupy burrows in shallow, tropical waters worldwide. Most of these animals inhabit structurally complex, benthic environments with an abundance of visual features that are regularly observed, including conspecifics, predators, prey, and landmarks for use in navigation. While these animals are capable of learning and discriminating color and polarization, it is unknown what specific attributes of a visual object are important for its recognition. Here we show that mantis shrimp of the species *Neogonodactylus oerstedii* can learn the shape of a trained target. Furthermore, when the shape and color of a target which they had been trained to identify were placed in conflict, *N. oerstedii* significantly chose the target of the trained shape over the target of the trained color. Thus, we conclude that the shape of a target is more important than its color for its recognition by *N. oerstedii*. Our findings suggest that the shapes of learned structures, such as landmarks or other animals, are important for *N. oerstedii* during object recognition.

## Introduction

Each species of animal living in a given space experiences its own distinct sensory world, known as its “umwelt” (von Uexküll, 1957/1934). The sensory structures responsible for an animal’s perception of its environment are metabolically taxing tissues that are often under strong selection pressures to permit the recognition of biologically relevant stimuli, while ignoring much of the available information an environment has to offer. Despite their complexity, the visual systems of stomatopod crustaceans should follow this generalization. Better known as mantis shrimp, these animals are renowned for their visual systems which in most species enable spatial and motion vision, color and multispectral UV vision, and linear and circular polarization receptivity (Cronin et al., 2014a). The compound eyes of many stomatopod species have a relatively high visual acuity; for instance, *Gonodactylus chiragra*, an animal typically about 8 cm in length, achieves a resolution of 0.8 cycles/degree (Marshall and Land, 1993). The ability of stomatopods to learn novel visual stimuli has been previously demonstrated with color, linear polarization, and circular polarization cues (Marshall et al., 1996; Marshall et al., 1999; Chiou et al., 2008; Thoen et al., 2014). Taken together, it is clear that visual information is an important part of a stomatopod’s sensory experience and likely critical for its survival.

Mantis shrimp mostly reside in shallow tropical marine waters worldwide. These locations offer some of the most structurally complex and colorful environments on Earth, and therefore contain many visual features. In these environments, mantis shrimp typically occupy small holes or crevices in the marine substrate for use as burrows, where they reside concealed for most of the day. Mantis shrimp consume a variety of prey (deVries et al., 2016), many of which are brightly colored, and they use colored signals to communicate with one another (Hazlett, 1979; Cheroske et al., 2009; Chiou et al., 2010; Franklin et al., 2019). Furthermore, mantis shrimp of the species *Neogonodactylus oerstedii* exhibit impressive navigational abilities when returning to their burrows from foraging excursions. These animals use landmarks, if available, in parallel with path integration to quickly pinpoint the location of their burrows (Patel and Cronin, 2020a,b,c). The benthic habitats *N. oerstedii* occupy are abundant with potential visually informative features including sponges, coral, rock, and aquatic vegetation: structures of distinct shapes and colors.

Since color may be informative in many aspects of a mantis shrimp’s life and since these animals use landmarks for navigation when available, this raises the question of what makes an object salient to a mantis shrimp for recognizing it. Considering that mantis shrimp have reasonably acute visual systems and are known to possess color vision, we were interested in determining whether *N. oerstedii* learns to recognize a visual target using its shape and/or its color.

## Results

### *Neogonodactylus oerstedii* learns to identify a specific visual target over time

*N. oerstedii* individuals (n=78) were trained to one of four targets of a specific color and shape combination (either a red rectangle, red triangle, green rectangle, or green triangle) using a paired food reward in a dichotomous choice y-maze (Fig 1). Since stomatopods in previous behavioral experiments successfully learned to discriminate red and green colored targets (Marshall et al., 1996), targets of these colors were chosen for the current study. The target of the alternate color and shape of the trained target was placed in the other arm of the y-maze and was associated with no reward (for example, a rewarded red triangle was paired with an unrewarded green rectangle). Animals on average responded in this situation (i.e. made a choice) approximately half of the time (Fig. 2A). From these choices, animals learned to associate food with their respective trained targets over time (P < 0.05; Fig. 2B). Of the 78 stomatopods that were trained, 20 individuals reached the criteria set to progress to the testing procedure (see the Methods section for the criteria).

**Figure 1.**
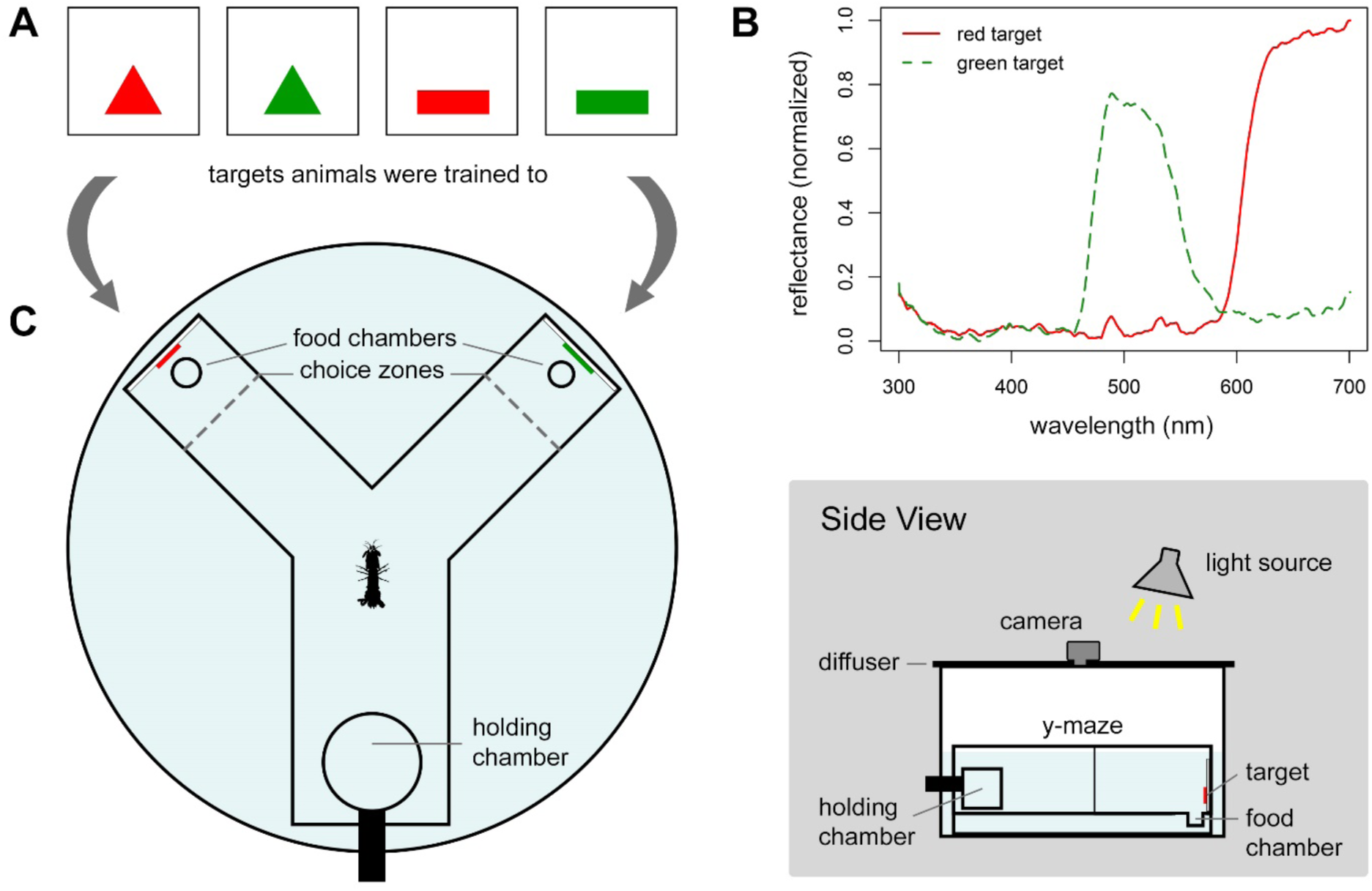
Layout of experimental setup. **(A)** The four targets used during the experiment: a red triangle, a green triangle, a red rectangle, and a green rectangle. **(B)** Averaged reflectance spectra (300 to 700 nm) of the red targets (solid red line) and green targets (dashed green line). **(C)** A y-maze was placed in a cylindrical tank with an incandescent light source centered above it. A diffusing filter was placed above the arena. The filter had a centered hole, where the lens of a camera was fit to record each trial. The y-maze contained an entrance arm and two choice arms oriented 90 degrees from one another. A cylindrical holding chamber was centered at the end of the entrance arm. At the end of each choice arm laid a hole set below the floor of the y-maze. A food reward could have been placed in either hole. One of the targets in (A) was placed at the end of each choice arm as indicated.

**Figure 2.**
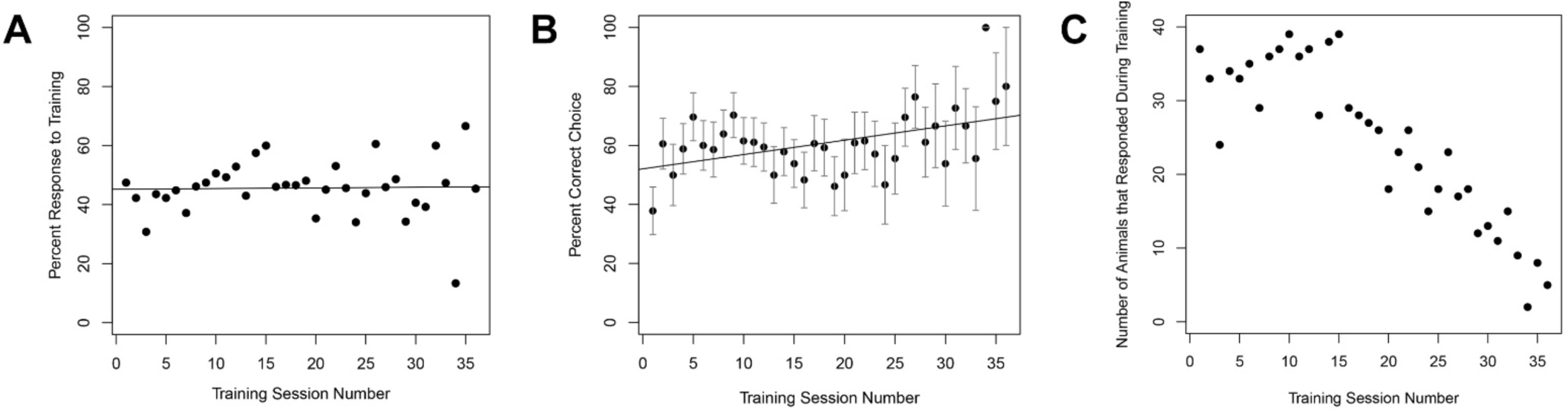
Training Results. **(A)** *Neogonodactylus oerstedii* responded to the training procedure approximately half of the time. **(B)** *N. oerstedii* associated food with their respective trained targets over time. Each point represents the percentage of animals who correctly chose the target they were training to from all animals who made choices during that training session. Error bars represent standard errors of the means. P = 0.034, r = 0.35. **(C)** Sample size per point in (B). The number of animals undergoing training decreased over time because animals either progressed to the testing procedure or died during the course of the study.

### *Neogonodactylus oerstedii* recognized the trained target by its shape, not its color

Once animals reached the performance criteria to enter the testing phase, they were tested in three separate procedures: a shape recognition test, a color recognition test, and a conflicting cue test. During all testing procedures, food was not paired with a target.

During the shape recognition test, both arms contained targets of the color an animal had been trained to, but the target in each arm was of a different shape (e.g. a red triangle paired with a red rectangle). In this experiment, individuals significantly chose the arm with the shape to which they had been trained, indicating that they recognized the shape of their trained target (P < 0.05, Z = 1.976, n = 19; Fig. 3).

**Figure 3.**
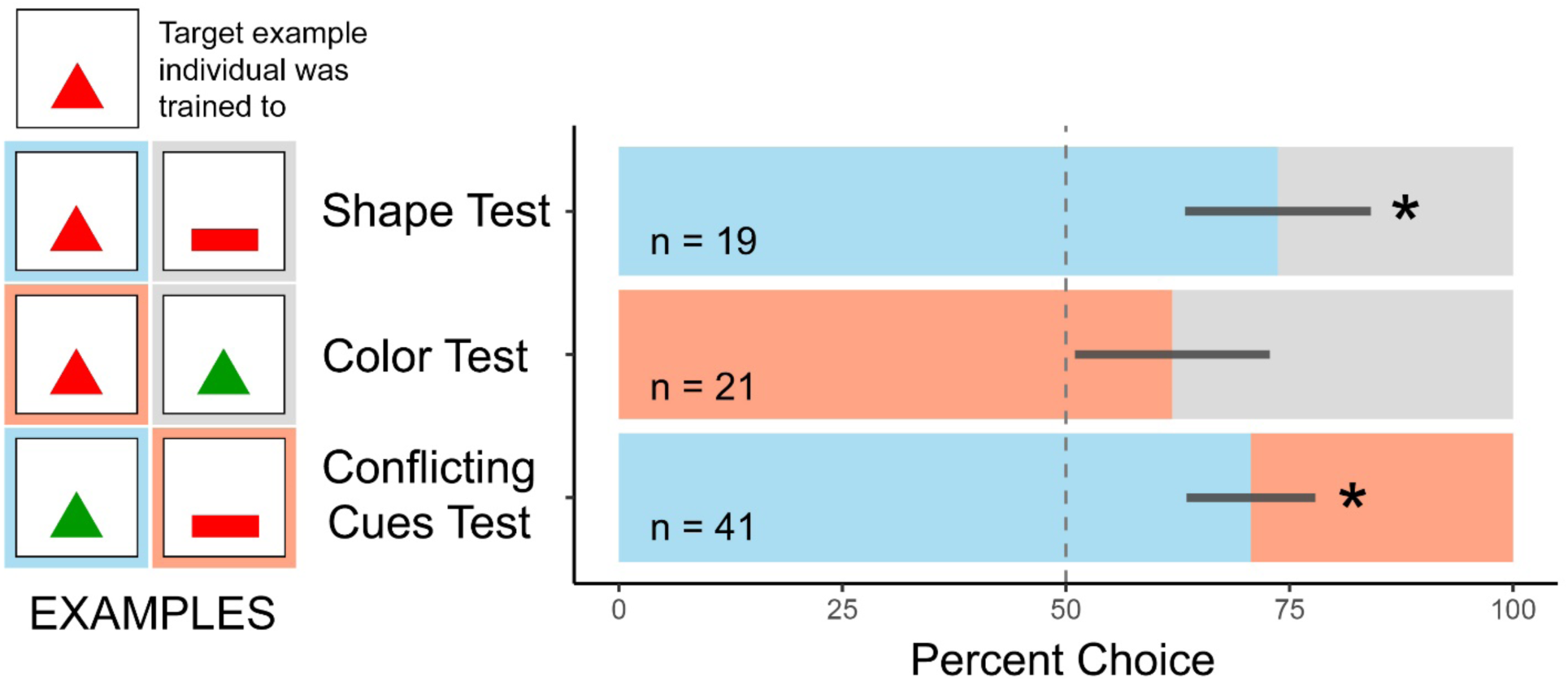
*Neogonodactylus oerstedii* recognized the target by its shape, not its color. Blue and red bars represent proportions of choices during testing that were for the target of the correct shape and color, respectively. Grey bars represent proportions of choices during testing that were for the incorrect target. Dark grey lines represent standard errors of the means. Asterisks (*) indicate a significant deviation from a random choice proportion (P < 0.05). The vertical dashed line marks a 50% proportion of choices (i.e. a random choice proportion). Examples of targets placed in either arm of each test for an individual that was trained to associate food with a red triangle are found on the left of each bar.

During the color recognition test, both arms contained targets of the shape they were trained to but the color of the target differed per arm (e.g. a red triangle paired with a green triangle). During this task, stomatopods more often than not chose the arm with the same color target that they were trained to; however, this relationship was not significantly different from a random choice distribution (P > 0.1, Z = 0.934, n = 21; Fig 3).

During the conflicting cue test, one arm contained a target with the same shape but opposite color to the target to which they were trained while the other arm had a target with the same color but alternate shape to the trained target (e.g. a green triangle paired with a red rectangle). Consistent with the results from the other tests, stomatopods significantly chose the arm with the trained shape over the arm with the trained color, implying that the shape of the trained target was more important than the target’s color to *N. oerstedii* during recognition (P < 0.05, Z = 1.927, n = 41; Fig. 3).

## Discussion

Our study is the first to demonstrate that mantis shrimp are able to recognize distinct shapes. We found that mantis shrimp ranked the shape of an object higher than its color when recognizing it (Fig. 3). Since mantis shrimp use landmarks during navigation (Patel and Cronin, 2020c), the findings in our study suggest that the shape of a landmark may be more important than its color when being identified by a mantis shrimp during navigation. Similarly, the shapes of prey, predators, and of body structures used in signaling may be critical for recognition and for generating appropriate behavioral responses independent of their roles in navigation.

Identifying an object by its shape might be more effective than recognizing its color when viewed underwater. In water, contrast attenuates with distance and depth due to the absorption and scattering of light. This is especially true for color information underwater, where the spectral range of incoming daylight or of an object’s color is rapidly trimmed to primarily blue light with increasing distance and depth (Cronin et al., 2014b). Because of this, luminosity contrast persists farther than color contrast in water, and the colors of objects vary with the distance and the depth of viewing while their shapes remain unchanged. Therefore, the shapes of objects may be more reliable cues to their identity than their colors when viewed by mantis shrimp in ecologically relevant situations.

Many animals use the edges of objects for recognition, including humans (Shapley and Tolhurst, 1973) and honeybees (Lehrer et al., 1990), so it is reasonable to hypothesize that mantis shrimps do the same. Shape recognition is likely to be critically important to mantis shrimp when they are recognizing landmarks, which they use to locate their home burrow during navigation (Patel and Cronin, 2020c). In other arthropods, landmark navigation involves retinal image matching, where the field of view seen while navigating is matched to a stored retinal “snapshot” of the view of their goal (Cartwright and Collett, 1983; Akesson and Wehner, 2007). During these tasks, the edges of landmarks appear to be important for image matching and distance estimations (Cartwright and Collett, 1983; Harris et al., 2007). Similarly, the overall panorama, especially the upper edge of the horizon, is used for orientation in some arthropods (Collett, 1996; Luschi et al, 1997; Fukushi, 2001; Graham and Cheng, 2009, Wystrach et al., 2011). Therefore, edge detection of objects may be critical during navigation as well as for other aspects of a mantis shrimp’s life, such as signal recognition, food identification, and recognition of predatory threats.

Conditioning experiments with other animals have demonstrated that multiple redundant cues can compete during associative learning, allowing one cue to overshadow the learning of another one (Couvillon et al., 1983; Menzel, 1990). In our experiments, we combined shape and color during associative learning. The apparent failure of our experimental animals to choose a target on the basis of color in our experiments suggests that shape was a more relevant cue in the task we gave the mantis shrimp, and therefore may have overshadowed the learning of the color of the target. On the other hand, while color recognition did not reach significance in these experiments, animals selected the color to which they had been trained over 60% of the time in our color recognition test and have significantly learned to recognize and discriminate color in the past (Marshall et al., 1996). These results suggest that further examination of the relevance of color in object recognition is warranted. For example, when shapes are similar, color may become more important in discriminating them.

Most mantis shrimp possess fabulously elaborate color vison systems. While color in our study did not seem to be important for object recognition, mantis shrimp are likely to use color discrimination for other specific tasks. Many mantis shrimp have colorful body surfaces, some of which are used for signaling (Hazlett, 1979; Cheroske et al., 2009; Chiou et al., 2010; Franklin et al., 2019). Due to mantis shrimps’ powerful weaponry and aggressive territoriality, signaling intent may be an important way to circumvent a potentially fatal encounter. Many mantis shrimp species possess colorful signals on the inner sides of their raptorial appendages termed meral spots. The colors of these spots often are distinct in coexisting species. Since multiple stomatopod species are often found occupying the same reef patches, the color of signals such as these meral spots might be useful for species recognition when identifying conspecifics. Color vision might also have other functions for mantis shrimp such as contrast enhancement when hunting and/or avoiding predators at shallow depths (Cronin et al., 2014c). Carl von Hess (1913) and Karl von Frisch (1914), early researchers studying color vision in honeybees, disagreed about the abilities of these animals to discriminate color (even though across the Atlantic, experimentation by Charles Turner (1910) strongly suggested that honeybees possessed color vision). The disagreement arose because the researchers chose different behavioral contexts in their studies. We now know that bees use color for nest and flower identification (the contexts in which Turner and von Frisch tested color vision), not for escape runs toward light (von Hess’s approach; see Menzel & Backhaus, 1989). Similarly, a mantis shrimp’s reliance on color vision surely differs depending on the contextually varied situations it encounters.

## Materials and Methods

### Animal Care

Individual *Neogonodactylus oerstedii* collected in the Florida Keys, USA were shipped to the University of Maryland Baltimore County (UMBC). Animals were housed individually in 30 parts per thousand (ppt) sea water at room temperature under a 12:12 light:dark cycle. Animals were fed whiteleg shrimp, *Litopenaeus vannamei*, once per week when food was not acquired during training sessions. 78 individuals (31 males and 47 females) that survived over four weeks in captivity were used for the study. Testing data were collected from 22 individuals (9 males and 13 females). All individuals were between 30 and 70 mm long from the rostrum to the tip of the telson.

### Experimental Apparatus

A y-maze consisting of an entrance arm and two choice arms oriented 90 degrees from one another was constructed out of while acrylic sheets (Figure 1). The end of each arm of the y-maze had a hole in the floor, hidden when viewed from a distance. A food reward was placed in either of these holes. The y-maze was placed in a cylindrical tank with an incandescent light source (Sylvania SPOT-GRO® 65W) centered above it. A diffusing filter was rested on the top of arena below the light source. The filter had a centered hole, where the lens of a small video camera was fit to record each trial. Trials were observed from the screen of this camera. Flat targets made of colored, transparent plastic placed on a solid white background were placed at the end of each choice arm. Four targets were used during the experiment: a red rectangle, a green rectangle, a red triangle, and a green triangle. The rectangle and triangle had an angular size (width x height) of 12° x 4° and 9.3° x 7.8° when viewed from the entrance to the choice arm, respectively. Targets were constructed from transparent plastic colored filters cemented to opaque white acrylic sheets (Figures 24D, 25). A cylindrical holding chamber was centered at the far end of the entrance arm. The holding chamber was designed to be rotated on its side by a researcher, allowing an animal placed inside the chamber access to the rest of the y-maze.

### Spectrometry

Reflectance measurements of the colored targets were taken in a dark room using an Ocean Optics USB2000 spectrometer connected to a 3 m long, 400 µm diameter, fiber-optic cable. Reflectances were measured from 300 to 700 nm relative to a “Spectralon” white standard using a PX-2 pulsed xenon light source.

### Experimental Procedures

#### Training

Each *N. oerstedii* individual was randomly assigned to be trained to one of the four target color and shape combinations described above. During training trials, the focal target (ex. red triangle) was placed at the end of a randomly chosen arm with food in the chamber at its end as a reward. The target of opposite shape and color (ex. green rectangle) was placed without food at the end of the other arm. A stomatopod was placed in the holding chamber before a trial and allowed five minutes to adjust to its surroundings. After this time, the holding chamber was turned, allowing the animal to enter the arena, initiating the experiment. Once a stomatopod entered the arena, the first choice arm it traveled down was noted once it entered the choice zone of the arm, two-thirds of the length of the arm. Once the food had been found, the experimental animal was allowed five minutes to eat as a reward before being removed from the arena. If the food was not found within 10 minutes, the animal was removed from the arena. Each animal experienced the training procedure twice per week. After each individual training session, the water in the arena was mixed to prevent olfactory cues from influencing the choice of subsequent training sessions.

At the end of each week, the percentage of correct choices each individual made since the start of training was calculated. Individuals entered the testing phase when they had made a correct choice 80% (or greater) of the time during training trials over the previous four weeks, in combination with a 50% (or greater) response rate during that time. Individuals were required to have been trained for at least one month (eight training trials) to be considered for testing.

#### Testing

Of the 78 animals that were trained, a total of 20 animals achieved the training criterion and moved on to the testing phase. The procedure of the testing phases was identical to that of the training phase except that no food reward was offered during testing sessions. Trained stomatopods were subjected to three distinct tests: a shape recognition test, a color recognition test, and a conflicting cues test (Fig. 3). Individuals experienced these tests in a randomized order. Two training sessions were administered between tests to facilitate reward seeking between tests.

1. The Shape Recognition Test In order to test if *N. oerstedii* could distinguish the shape of the trained target, the cue of the same shape and color as what the individual was trained to was placed at the end of one arm of the Y-maze (ex. red triangle). The cue of the opposite shape and the same color of what was trained to was placed at the end of the other arm (ex. red rectangle). A correct choice was recorded if the stomatopod chose the arm displaying the cue with the trained color and shape.
2. The Color Recognition Test In order to test if *N. oerstedii* could distinguish the color of the trained target, the cue of the same shape and color as what the individual was trained to was placed at the end of one arm of the Y-maze (ex. red triangle). The cue of the same shape and the opposite color of what was trained to was placed at the end of the other arm (ex. green triangle). A correct choice was recorded if the stomatopod chose the arm displaying the cue with the trained color and shape.
3. The Conflicting Cues Test In order to test if *N. oerstedii* relied more on the shape or color of a target when recognizing it, the cue of the same shape and opposite color as what the individual was trained to was placed at the end of one arm of the Y-maze (ex. green triangle). The cue of the opposite shape and the same color of what the animal was trained to was placed at the end of the other arm (ex. red rectangle). Neither cue was of the identical shape and color combination to the one to which the animal was trained to recognize.

### Statistical Analyses

All statistical analyses were run on R (v3.3.1, R Core Development Team 2016) with the “car”, “glmer”, and “lme4” plugins.

A Pearson’s correlation test was used to compare the proportion of correct choices made during training sessions over time.

Generalized linear mixed modelling was used to analyze the data for each of the three tests. Our models used animal choices during testing as the variable of interest, specifying a binomial error distribution (link function “logit”). Since individual stomatopods were tested more than once, the models for each test included individual ID as a random term. Since we used both males and females for our study, sex was also included as a random term for our full models; however, since sex did not significantly increase the explanatory power of our models, it was removed from our final models. Individual ID did not significantly increase the explanatory power of our models, but was left in the final models to account for repeated measures.

## Acknowledgements

We thank N.S. Roberts for discussions about appropriate statistical analyses for the study.

## Author Contributions

R.N.P. designed the research, analyzed the data, and prepared the manuscript. V.K, L.A., H.D., T.G., S.P, O.P.R., A.S, and B.S collected the data. V.K. oversaw data collection and managed the data. T.W.C. provided guidance and research support.

## Data and Materials Availability

The data that support the findings of this study are available from the corresponding author upon reasonable request. Correspondence and requests for materials should be addressed to R.N.P.

**Table 1:** Summary of generalized linear mixed modelling outcomes for all experimental tests.

| Experiment | P-value | Z | n (tests) |
| --- | --- | --- | --- |
| Shape Recognition Test (Chose correct shape) | 0.0481 | 1.976 | 19 |
| Color Recognition Test (Chose correct color) | 0.35 | 0.934 | 21 |
| Conflicting Cues Test (Chose correct shape) | 0.054 | 1.927 | 41 |
| Degree to which repeated measures explain results | P-value | X<sup>2</sup> | n (individuals) |
| Shape Recognition Test* | 1 | 0 | 12 |
| Color Recognition Test* | 0.432 | 0.6187 | 13 |
| Conflicting Cues Test* | 0.227 | 1.4577 | 15 |

\*Repeated measures do not significantly explain results.

